# Identifying, Prioritizing, and Visualizing Functional Promoter SNVs with the Recurrence-agnostic *REMIND-Cancer Pipeline* and *pSNV Hunter*

**DOI:** 10.1101/2024.09.10.612212

**Authors:** Nicholas Abad, Irina Glas, Chen Hong, Yoann Pageaud, Barbara Hutter, Benedikt Brors, Cindy Körner, Lars Feuerbach

## Abstract

Cancer is a heterogeneous disease that arises due to mutations that drive cancer progression. However, the identification of these functional mutations has typically focused only on protein-coding DNA. Among non-coding mutations, only a few have been clearly associated with cancer. We hypothesize that this gap in discovery is partly due to the limitations of current methods requiring high recurrence of mutations. To support candidate selection for experimental validation of lowly recurrent and singleton promoter mutations, new computational approaches for the integrated analysis of multi-omics data are required.

To address this challenge, the *REMIND-Cancer Pipeline* leverages whole-genome sequencing and RNA-Seq data to extract and prioritize functional promoter mutations, regardless of their recurrence status. Subsequently, *pSNV Hunter* aggregates and visualizes comprehensive information for each candidate. We demonstrate the functionality of both tools by applying it to the PCAWG dataset. This workflow successfully identified and prioritized known highly-recurrent mutations, as well as, novel singletons and lowly recurrent candidates. Hence, the output of our workflow directly supports hypothesis generation for subsequent experimental validation to overcome limitations of recurrence-based approaches.

## INTRODUCTION

Cancer is a heterogeneous disease that arises from genetic mutations that confer a growth advantage to the cancer cells by modifying the activity of cellular pathways. The most common type of somatic mutations are single nucleotide variants (SNVs) [1,2]. Historically, SNVs and their impact have mostly been studied within the coding region of DNA [3,4]. However, whole genome sequencing (WGS) enables a comprehensive view of an entire cancer genome, including the non-coding region that constitutes 98% of human DNA [5].

The investigation of the non-coding genome in the context of cancer led to the landmark discovery of two hotspot SNVs within the promoter region of telomerase reverse transcriptase (*TERT*), namely *TERT*_*C228T*_ and *TERT*_*C250T*_ [6,7]. These highly-recurrent promoter SNVs (pSNVs) were shown to lead to an upregulation of *TERT* in multiple cancer types by creating new transcription factor binding sites (TFBSs) for ETS-family transcription factors (TFs). Since their discoveries, only a select few other functional pSNVs, such as *RALY*_*C927T*_ [8] and *CDC20*_*G529A*_ [9,10], have been discussed in the literature. These discoveries were enabled through computational tools such as MutSigCV [11], oncodriveFML [12], FunSeq2 [4], LARVA [13] and ncDriver [14].

These methods generally follow a four-step process: (1) calculate an *observed* mutation rate for a region of interest (e.g. promoter region), (2) estimate an *expected* mutation rate (i.e. background model) potentially based on cancer type among other features, (3) compare these two distributions, and (4) determine whether the observed rate is statistically different than expected. However, this methodology inherently depends on one important factor: recurrence. Previous studies have shown that in order to achieve at least 90% statistical power using these recurrence-based mutational burden tests, a mutation needs to be present in at least 1% to 25% of samples depending on cohort size and the observed cohort mutation rate [3]. This therefore implies that singletons (i.e. SNVs that only occur in a single sample) and lowly-recurrent pSNVs cannot be differentiated from random noise using this approach [15].

To also enable the detection of these singletons and lowly recurrent pSNVs, we created the Regulatory Mutation Identification ‘N’ Description in Cancer (REMIND-Cancer) pipeline, which incorporates background knowledge about TFBS sequences and gene expression to incorporate additional features. Hence, it is a recurrence-agnostic pipeline aimed at identifying and prioritizing functional pSNVs that affect TFBSs and increase the expression of its corresponding gene. To assess its functionality, we thus applied this pipeline to the Pan-Cancer Analysis of Whole Genomes (PCAWG) [16] dataset, which comprises 2,413 samples and 19,401,901 mutations that span 43 distinct cohorts. To assist in the manual inspection phase following the filtering and ranking steps, a comprehensive data aggregation and interactive visualization tool, *pSNV Hunter*, was developed. Both tools are freely available.

## METHODOLOGY

The REMIND-Cancer pipeline follows a filtering-ranking-inspection paradigm in which somatic pSNVs identified in tumor WGS samples are (1) filtered to attempt to focus only on putative driver mutations, (2) scored and ranked based on genomic, transcriptomic, and annotation features, and (3) manually inspected to remove artifacts and ensure relevance for any downstream analysis.

### PCAWG Dataset

Both the WGS (.vcf file format) and RNA-Seq (.tsv file format) data used in this study were from the publicly-available PCAWG dataset [16]. To match a sample’s WGS to their RNA-Seq data, the specimen’s submission ID (WGS) was matched to its corresponding submitter bundle ID (RNA-Seq). Most WGS cancer types had corresponding RNA-Seq data available (Supplementary Figure 1). In the case of the early onset prostate cancer (EOPC-DE) cohort, only WGS data was publicly available. However, we integrated this with its separately-published RNA-Seq data [17]. In total, this dataset comprises 2,413 samples and 19,401,901 SNVs.

### Promoter Filter

The promoter region was defined as ranging from 1,000 base pairs upstream to 500 base pairs downstream of the transcription start site (TSS). To be consistent with the PCAWG [16] project, all coordinates are based on the GRCh37 (hg19) genome assembly and the GENCODE annotation V19 [18]. Only the canonical transcripts were considered (Supplementary Table 1). Utilizing each samples’ somatic SNV file in Variant Call Format (VCF) [19], only mutations within the promoter were kept for further analysis.

### Gene Expression Filter

To identify upregulated genes, we utilized the RNA-Seq-based gene expression data available for 950 samples and therefore disregarded pSNVs from samples lacking RNA-Seq data. The gene expression, measured in fragments per kilobase of transcript per million mapped reads (FPKM), of the gene adjacent to the mutated promoter was z-score normalized relative to samples within its subcohort. A z-score greater than 0 then indicated an above-average expression and the corresponding pSNVs were retained.

### TFBS Motif and Expression Filter

Mutations that impacted the binding motif of transcription factors were identified within this filter. Utilizing the Find Individual Motif Occurrences (FIMO) [20] tool from the MEME Suite toolkit [21] along with the JASPAR2020 [22] database of curated transcription factors, the hg19 reference DNA sequence of +/-10 bp around every pSNV underwent scanning by FIMO against every TF motif in JASPAR2020. Using the positional weight matrix of the TF and its associated sequence context, FIMO calculated a *statistical binding affinity score*, which indicates the likelihood of observing the specific motif. To assess the impact of a mutation, the ratio (denoted as S(TFBS)) between the binding affinity scores for the mutant over the wild type allele was calculated as a quantitative measure indicating the impact of mutations on the binding motif of TFs. Motifs with S(TFBS)>11 were defined as a created TFBS motif and S(TFBS)<0.09 as a destroyed TFBS motif. Mutations were kept if at least one predicted TF fulfilled one of these criteria. pSNVs were retained if it altered at least one TFBS for which the matching TF had a FPKM > 0 based on the RNA seq data.

### Recurrence, CGC, and Chromatin Accessibility Annotations

Additional pSNV features were annotated for computing the prioritization score and subsequent ranking. Among these features, recurrence was determined by utilizing all available PCAWG WGS data, irrespective of RNA-Seq availability. Recurrence was determined by creating an index of all pSNVs based on their chromosome, position, and nucleotide change using a Python dictionary and by querying the number of entries for each locus to obtain a recurrence statistic for each sample-specific pSNV.

Furthermore, pSNVs matching to genes listed as one of the 746 entries in the Cancer Gene Census (CGC) database [23] as of 11 November 2020 were labeled as CGC “True” (Supplementary Table 2). All CGC tiers of evidence were equally considered for the prioritization scoring.

To annotate the presence of open chromatin overlapping with the pSNV, their positions were intersected with the ChromHMM [24] annotation as found in the ‘hg19_genome_100_segments’ file which was last modified on 5 December 2022. These chromatin-state annotations were based on their universal full-stack model, which was created by training a hidden markov based model on 1,000+ datasets within 100+ cell types. Any pSNV labeled as being within the regions of a bivalent promoter (i.e. BivProm1-4) or containing one of the histone modifications H3K4me1-3 (i.e. PromF1-5) were labeled as having open chromatin (Supplementary Table 3).

### Candidate Mutation Prioritization

To prioritize the most promising candidate pSNVs, an empirically calibrated weighted-sum scoring function was used to compute a prioritization score for each sample-specific pSNV. The function favorably scored pSNVs with strong upregulation, in regions of open chromatin, with high allele frequency indicating clonality and from samples with high tumor purity. Further, recurrence in other samples and association to known cancer genes increased the score and the mutation of multiple TFBSs was acknowledged.

The specific genomic, transcriptomic, and annotation features that were used and their corresponding weights are summarized in Supplementary Table 4. The default calibration of the weights prioritized specificity over sensitivity, though these weights, thresholds, and maximum contributions can be altered by the user. Briefly, the number of created or destroyed TFBSs is weighted by 2, both having a maximum contribution of 6 to the overall prioritization score (e.g. prioritization score contribution is *min(6, number of created TFBSs* × *2)*). The number of other samples harboring the pSNV (recurrence) is weighted by 5 with a maximum contribution of 25. If a sample’s tumor purity is greater than 0.25, a contribution of 10 is added. If a sample’s variant allele frequency exceeds 0.3, a similar contribution of 10 is added. The normalized expression of the pSNV’s corresponding gene was also considered by weighting the z-score by 5 with a maximum contribution of 25 to the overall prioritization score. Lastly, a pSNV residing in a location of open chromatin, according to ChromHMM, received an additional 20 points whereas a pSNV affecting a known cancer gene, according to the CGC database, receives 15 points.

More specifically, the prioritization score was computed using the following formula:

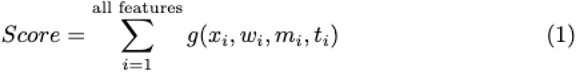

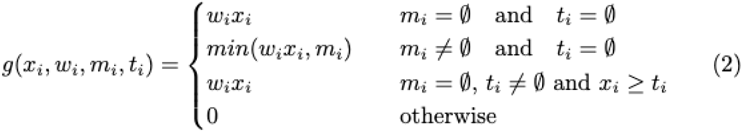

such that

- *Score* is the prioritization score of a pSNV
- *x*_*i*_ is feature *i*’s value
- *w*_*i*_ is feature *i*’s weight
- *m*_*i*_ is feature *i*’s maximum value
- *t*_*i*_ is feature *i*’s threshold

Of note, the scoring parameters can be adjusted to users’ preferences within the tool interface.

### Quality Control Measures

To manually assess the technical and potential biological validity of pSNVs, we employed two established tools, *DeepPileup* [3] and *GenomeTornadoPlot* [25], during the inspection phase. *DeepPileup* tests if the genomic positions of a SNV is prone to sequencing artifacts by analyzing pileups in the tumor and control samples of the full PCAWG dataset. Then, the percentage of samples per cohort in which more than two variants (2V+) are observed is visualized. Germline variants are identified as positions with a minor allele frequency > 25% in control samples. Remaining candidates in which the control sample reached a similar or higher 2V+ percentage than tumor samples in more than two cohorts were considered noisy. An example of a noisy genomic position can be seen in Supplementary Figure 2.

Further, for each pSNV-gene pair we employed the *GenomeTornadoPlot* package to test for an enrichment of focal high level amplifications in the remaining PCAWG cohort. This could be indicative of an alternative gene activation mechanism in other tumors and therefore imply convergent tumor evolution.

### pSNV Hunter

To assist in the inspection step of the REMIND-Cancer pipeline, we created *pSNV Hunter*, a multi-functional software tool that allows for the examination of individual pSNVs. Through an interactive dashboard that was created in Python using the Plotly Dash package, users can view plots displaying the gene expression of the adjacent gene corresponding to the pSNV, recurrence level, TF expression, and all plots provided by *DeepPileup* and *GenomeTornadoPlots*. Users can also view specific gene and TF functions provided by the National Center for Biotechnology Information (NCBI) [26] as well as leave comments for parallel analysis between multiple users. These main features can be seen in Figure 1B.

**Figure 1:**
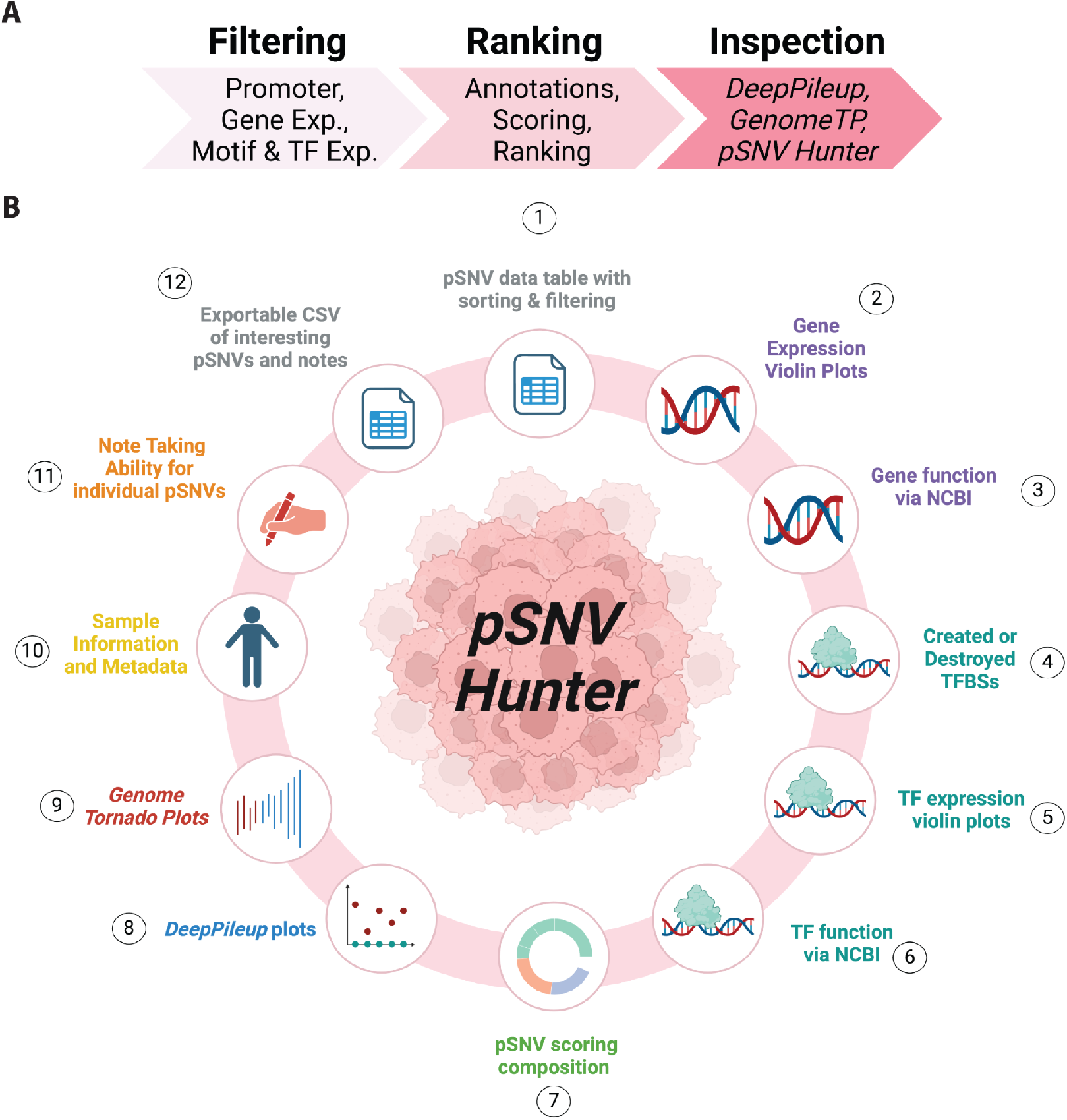
Schematic overview of the REMIND-Cancer approach and the main features of pSNV Hunter to assist users during the inspection phase. **(A)** Following a filtering-ranking-inspection paradigm, each stage consists of multiple steps. Within the filtering phase, three filters (promoter, gene expression and TFBS motif and TF expression) are applied. Secondly, within the ranking phase, annotations (e.g. recurrence, known cancer gene, open chromatin) are added and used to develop a prioritization score, which is thus used to rank sample-specific pSNVs. Lastly, two quality control measures (i.e. DeepPileup and GenomeTornadoPlots) are applied and can be viewed, along with all other pSNV information, using our comprehensive visualization tool pSNV Hunter. **(B)** An overview of the 12 most important features of pSNV Hunter. The different text colors represent distinct tabs within the tool itself: table-like sorting and filtering (grey; 1), gene-specific information (purple; 2-3), TF-specific information (teal; 4-6), pSNV scoring composition and breakdown (green; 7), DeepPileup QC plots (blue; 8), GenomeTornadoPlots plots (red; 9), general information and metadata about the sample (yellow; 10), note taking for users (orange; 11), and exporting of all notes and relevant information (grey; 12). Both figures were created with BioRender.com.

### Code Availability

Four code repositories are publicly-available via GitHub. (1) The REMIND-Cancer pipeline was developed using Python 3.10.11 and primarily utilized the following packages: Pandas (2.0.1) [27], NumPy (1.24.3) [28] and SciPy (1.11.2) [29]. This can be accessed at https://github.com/nicholas-abad/REMIND-Cancer. (2) *DeepPileup*, which was primarily developed in Python 3.10.11, can be found at https://github.com/nicholas-abad/deep-pileup-wrapper. (3) A wrapper (i.e. an abstraction of the original package) for *GenomeTornadoPlot* can be found at https://github.com/nicholas-abad/genome-tornado-plot-wrapper, which utilizes the original scripts found in https://github.com/chenhong-dkfz/GenomeTornadoPlot/. (4) *pSNV Hunter* is publicly-available at https://github.com/nicholas-abad/pSNV-hunter and can be used with not only REMIND-Cancer pipeline results but also with any VCF file.

## RESULTS AND DISCUSSION

### The REMIND-Cancer pipeline identifies and prioritizes previously-validated recurrent pSNVs while also highlighting numerous novel singletons

To identify putative functional pSNVs within the data from the PCAWG project, the REMIND-Cancer pipeline applied a series of filters. Initially, this dataset included 2,413 samples and 19,401,901 SNVs (Fig. 2A). By focusing only on samples with both WGS and RNA-Seq data, the dataset was refined to 950 samples from 25 distinct cohorts, corresponding to 10,981,205 SNVs for further analysis.

**Fig. 2:**
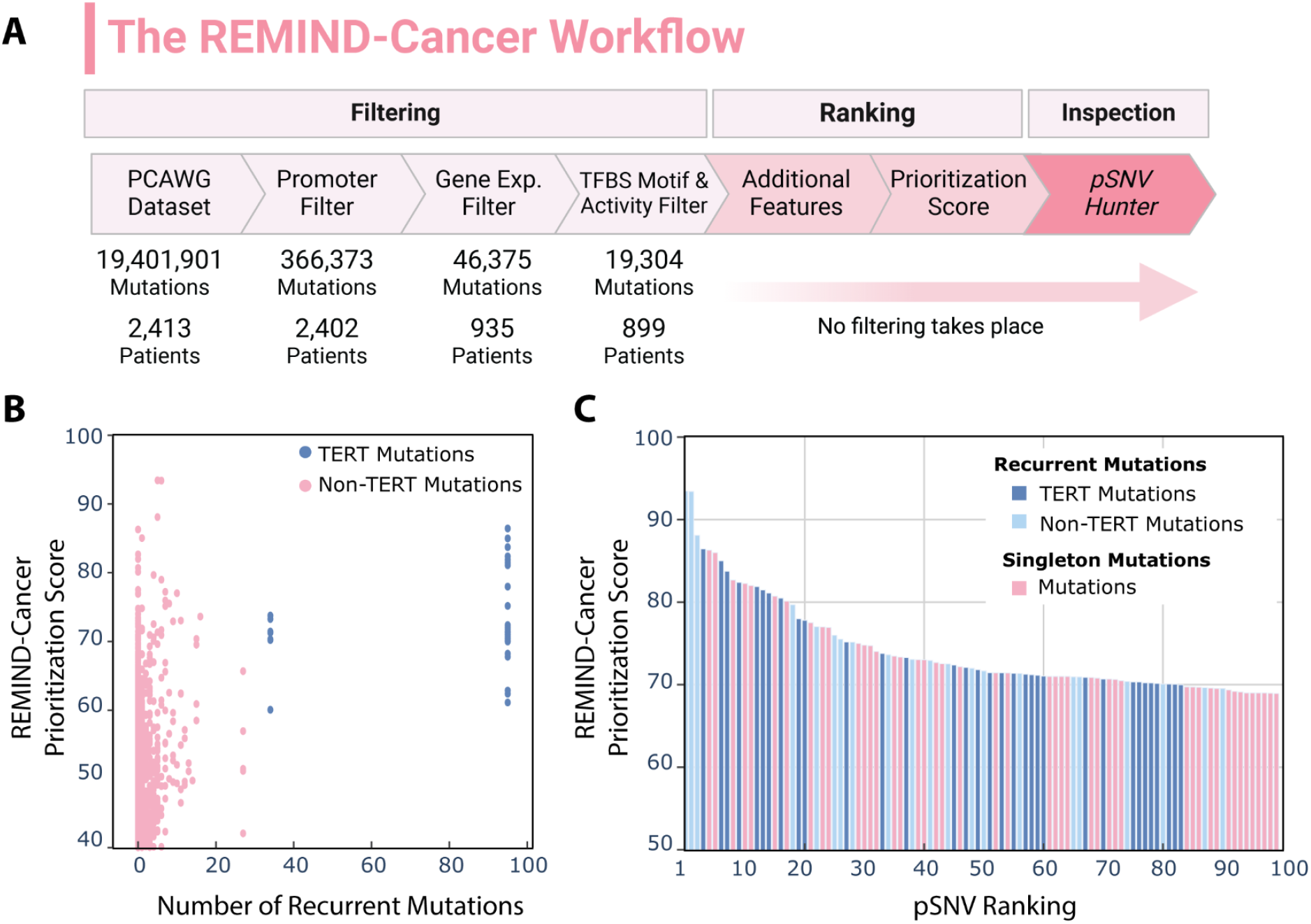
Characterization of the results of the REMIND-Cancer workflow. (a) An overview of the REMIND-Cancer workflow, which has four major components: filtering, ranking, inspection, and validation. Within each of these three components (e.g. filtering), the different steps (e.g. promoter filter) can be seen along with the number of mutations at the conclusion of this step (e.g. 366,373 mutations) and the number of patients that these mutations belong to (e.g. 2,402 patients) (b) For pSNVs that successfully passed the pipeline, the sample-specific recurrence level (mutation frequency minus 1; x-axis) is plotted against its REMIND-Cancer prioritization score (y-axis). Both TERT promoter mutations are depicted in dark blue and the recurrent non-TERT mutations in pink. (c) Distribution of the REMIND-Cancer Prioritization Score of the top 100 ranking pSNVs. Each bar represents a sample-specific pSNV and is colored by recurrence or singleton status and TERT association. The maximum score a pSNV can obtain is 117. Figure 2A was created with BioRender.com.

The application of our workflow identified 366,373 SNVs in promoter regions, i.e. pSNVs (Fig. 2A). By further applying the gene expression and TFBS motif and TF expression filters, the candidate set was reduced to 19,304 pSNVs, which occurred within 10,560 unique genes and 899 patient samples (Supplementary Table 5). These remaining pSNVs were then annotated according to mutational recurrence, tumor purity, variant allele frequency, inclusion of the matching gene in the Cancer Gene Census (CGC) database, and co-location with accessible chromatin, as detailed in the Methodology section. These features were used to compute an empirical prioritization score, which was then used for subsequent pSNV candidate ranking. Hereby, recurrent pSNVs obtained an individual score for each sample they occurred in due to individual gene expression values, expression of affected TFs, tumor purity estimates and variant allele frequency.

Many previously-implicated recurrent pSNVs, such as *TERT*_*C228T*,_ *TERT*_*C250T*,_ and *RALY*_*C927T*_, were identified and received high prioritization scores by our pipeline [6-8]. Specifically, of the 96 *TERT*_*C228T*_ sample-specific pSNVs within the entire dataset, 65 (68%) had corresponding RNA-Seq data and were thus eligible to pass the pipeline. As a result, 28 passed all filters and 26 of them were ranked within the top 100, indicating the capability of our pipeline to identify previously described pSNVs (Fig. 2B). Similarly, 7 of the 35 total (26 with RNA-Seq data) *TERT*_*C250T*_ pSNVs passed the pipeline as well as 4 of the 16 total (6 with RNA-Seq data) *RALY*_*C927T*_ pSNVs (Fig. 2B). Overall, *TERT*_*C228T*,_ *TERT*_*C250T*,_ and *RALY*_*C927T*_ made up 26, 7, and 2 instances, respectively, of the 54 recurrent mutations within the top 100 (Fig. 2C). Other than these three pSNVs, the only other recurrent mutations with multiple instances within the top 100 were *ANKRD53*_*G529A*_ and *SECISBP2*_*G357A*_ with exactly two occurrences each. Correspondingly, *ANKRD53*_*G529A*_ was observed 6 times within the entire dataset in comparison to the 7 instances of *SECISBP2*_*G357A*_.

Though recurrent mutations were prioritized, a majority of pSNVs (n = 18,337; 95%) passing the filtering steps were singletons and have not yet been implicated within the literature. Furthermore, 46 singletons were ranked within the top 100 (Fig. 2C). This is especially striking given the fact that recurrence was part of the score, thereby singletons lacking the corresponding scoring bonus. In principle, singletons hold the same potential of being biologically functional as recurrent mutations. Therefore, the identification and further analysis of this understudied category of mutations is necessary to expand our understanding of the complex gene regulatory networks within individual tumors.

### Investigating the known recurrent *CDC20* promoter mutation using *pSNV Hunter*

The previously-identified *CDC20*_*G529A*_ pSNV was recurrently found within the skin cutaneous melanoma (SKCM-US) cohort and prioritized within the top 5% of pSNVs by the REMIND-Cancer workflow (Fig. 3A). Upon closer inspection of the sample with the highest sample-specific pSNV score via *pSNV Hunter*, we observed an extremely high *CDC20* expression level (z-score of 2.2; Fig. 3B) as well as a high recurrence rate of 11. However, of the other samples harboring the *CDC20*_*G529A*_ pSNV, 9 were found within the Australian melanoma cohort (MELA-AU), which did not have RNA-Seq data available, whereas the other two samples corresponded to slightly negative expression z-scores (i.e. z-scores of -0.49 and -0.63).

**Fig. 3:**
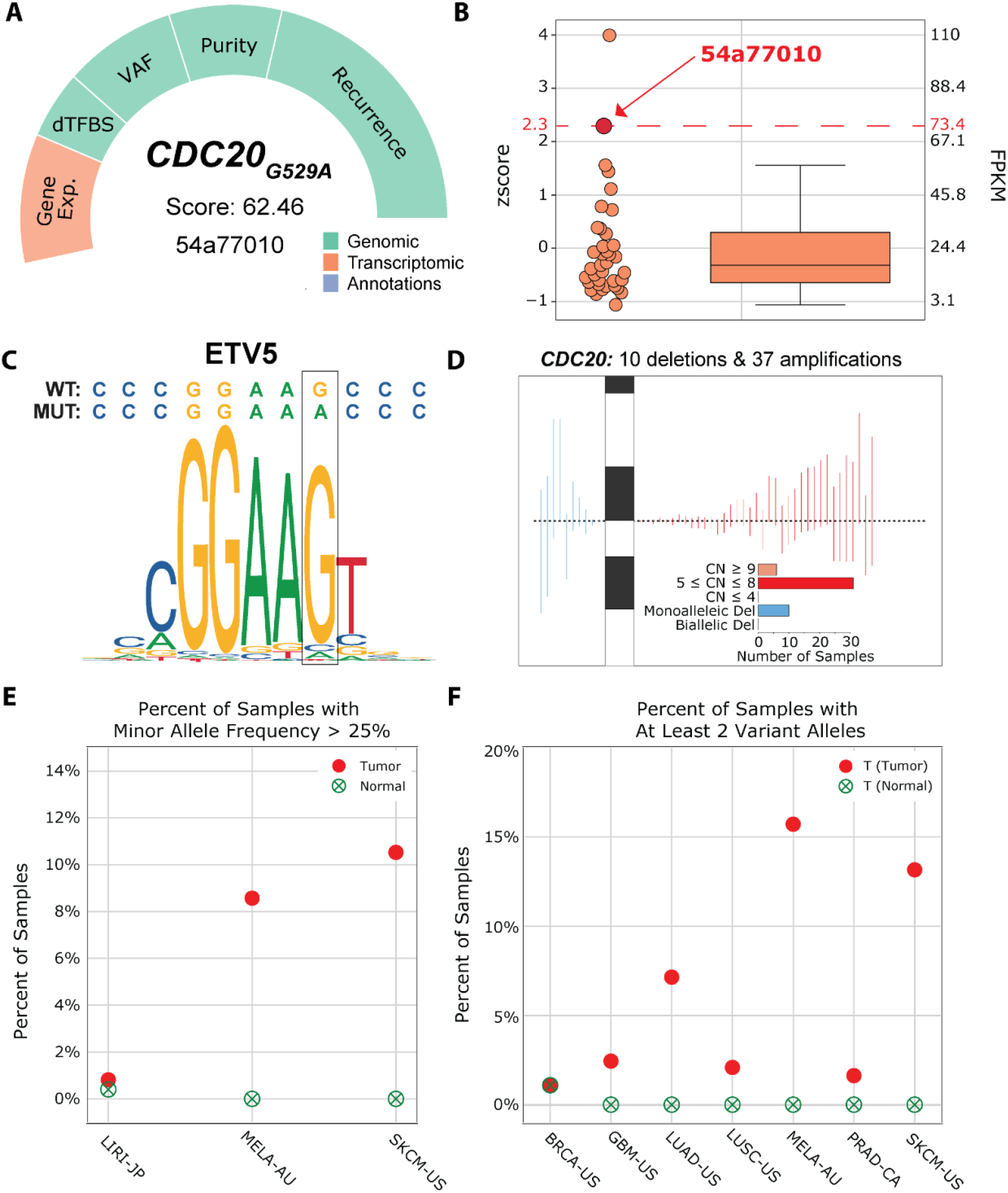
Overview of the high-ranking CDC20_G529A_ pSNV provided by pSNV Hunter. (A) Sample-specific scoring breakdown of the highest-ranking CDC20_G529A_ pSNV (B) The z-score (left y-axis) and FPKM (right y-axis) for all SKCM-US samples, highlighting the sample (54a77010) where this pSNV was detected (indicated by the red dot and line). (C) Logo plot demonstrating the creation of the ETV5 TFBS. (D) GenomeTornadoPlot showing the number of focal deletions and amplifications of CDC20. (E) DeepPileup plot of the percentage of samples with a minor allele frequency above 25% for both tumor and normal samples per cohort. (F) DeepPileup plot of the percentage of samples with at least two variant alleles for both tumor and normal samples per cohort.

Moreover, this pSNV was predicted to disrupt the binding of multiple TFs including that of the ETS-family TF ETV5 (Fig. 3C). This destruction was partially observed in a previous study in which the destruction of the TFBS of another ETS-family TF ELK4 that has the same TFBS motif as ETV5 was found to lead to the upregulation of *CDC20 in vitro* [9].

Furthermore, *pSNV Hunter* revealed that this location is amplified in 37 samples within the PCAWG dataset, of which a majority have a copy number (CN) above 4 (Fig. 3D). Observing two different activating mechanisms in the same gene in different samples is hinting at convergent tumor evolution. Interestingly, one of the samples harboring *CDC20*_*G529A*_ also has a copy number of 3. As the allele frequency of the mutation is 0.525 with a tumor purity of 0.74, this implies that the mutated allele within this sample is duplicated. This effect was also observed in another sample harboring the *CDC20*_*G529A*_ pSNV, which had an amplification (CN = 4) as well as a high relative expression value (zscore = 1.6).

Additionally, the *DeepPileup* plots also revealed by *pSNV Hunter* show technical and biological validity as signal is generally only observed within tumor samples and only in a single case within a normal sample of the BRCA-US cohort which may indicate a germline event (Fig. 3E and Fig. 3F).

Though not within the COSMIC database of known cancer genes [23], prior studies have investigated this specific pSNV as a potential biomarker [9,10], which points to an additional layer of investigation needed prior to any further downstream *in vitro* analysis. Altogether, this recurrent mutation was identified and prioritized by the REMIND-Cancer workflow and after extensive inspection via *pSNV Hunter*, we would suggest this pSNV as a promising target for functional validation *in vitro*, e.g. by promoter luciferase assays.

## CONCLUSION

Current recurrence-based methods require mutations being present in a large amount of samples in order to be identified. As a result, singletons and lowly-recurrent mutations are consistently overlooked in the list of known functional mutations, despite these mutations comprising a majority of genomic events. To address this limitation, we created the Regulatory Mutation Identification ‘N’ Description in Cancer (*REMIND-Cancer*), which identifies and prioritizes functional pSNVs through a recurrence-agnostic pipeline. When applied to the PCAWG dataset, this pipeline successfully identified and prioritized previously-known highly-recurrent pSNVs (e.g. *TERT*_*C228T*_,*TERT*_*C250T*_, and *CDC20*_*G529A*_) as well as novel singletons and lowly-recurrent mutations. Additionally, we created *pSNV Hunter*, a standalone data aggregation and visualization dashboard, to enable users to explore their findings in more detail as exemplified here for *CDC20*_*G529A*_. These results demonstrate that recurrence-agnostic approaches, such as the *REMIND-Cancer* pipeline, can effectively identify functional pSNVs, thereby potentially expanding the catalog of known functional mutations in cancer genomics. Collectively, our workflow overcomes the limitations of recurrence-based approaches and thereby directly supports hypothesis generation for subsequent experimental validation of novel pSNVs.

## AUTHOR’S CONTRIBUTIONS

Data Curation: N.A, L.F, I.G., C.H., Y.P.; Bioinformatic Analysis: N.A, L.F., I.G., C.K.; Writing: N.A., L.F., C.K.; Scientific Input and Editing: Y.P, B.H., B.B., C.P., I.G., D.W.

## COMPETING INTERESTS

There are no competing interests.

## ACKNOWLEDGEMENTS

We would like to thank Prof. Dr. Stefan Wiemann for his comments regarding the manuscript. NA has been supported by the Fritz Thyssen Stiftung’s REMIND Cancer grant and LF has been supported by the German Federal Ministry of Research and Education through the EOPC-Data Mining Grant (01KU1505A). We acknowledge the contributions of the many research networks across ICGC and TCGA who provided samples and data to the PCAWG Consortium. We thank the patients and their families for their participation in the individual ICGC and TCGA projects.

## SUPPLEMENTARY TABLES

1. GENCODE v19 promoter / TSS locations
2. CGC list used
3. ChromHMM regions
4. Weights/thresholds for scoring
5. Results of top 1,000 mutations (EOPC-DE included)

## SUPPLEMENTARY FIGURES

**Supplementary Fig. 1:**
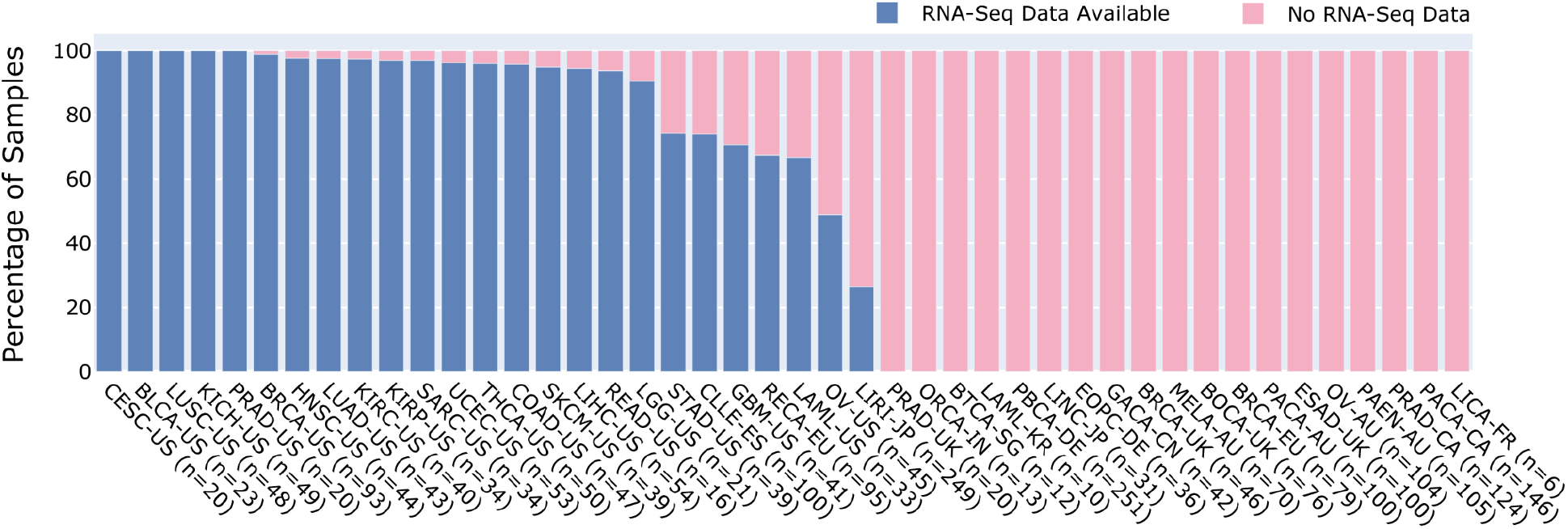
A bar plot representing the percentage of patients in each cohort that also have RNA-Seq data available within the PCAWG dataset. A value of 100% (i.e. CESC-US, BLCA-US, etc.) implies that every WGS sample has a corresponding RNA-Seq file. A majority of cohorts (i.e. LICA-FR, PACA-CA, etc.) did not have any samples with corresponding RNA-Seq data available and therefore did not pass the entirety of the REMIND-Cancer pipeline.

**Supplementary Fig. 2:**
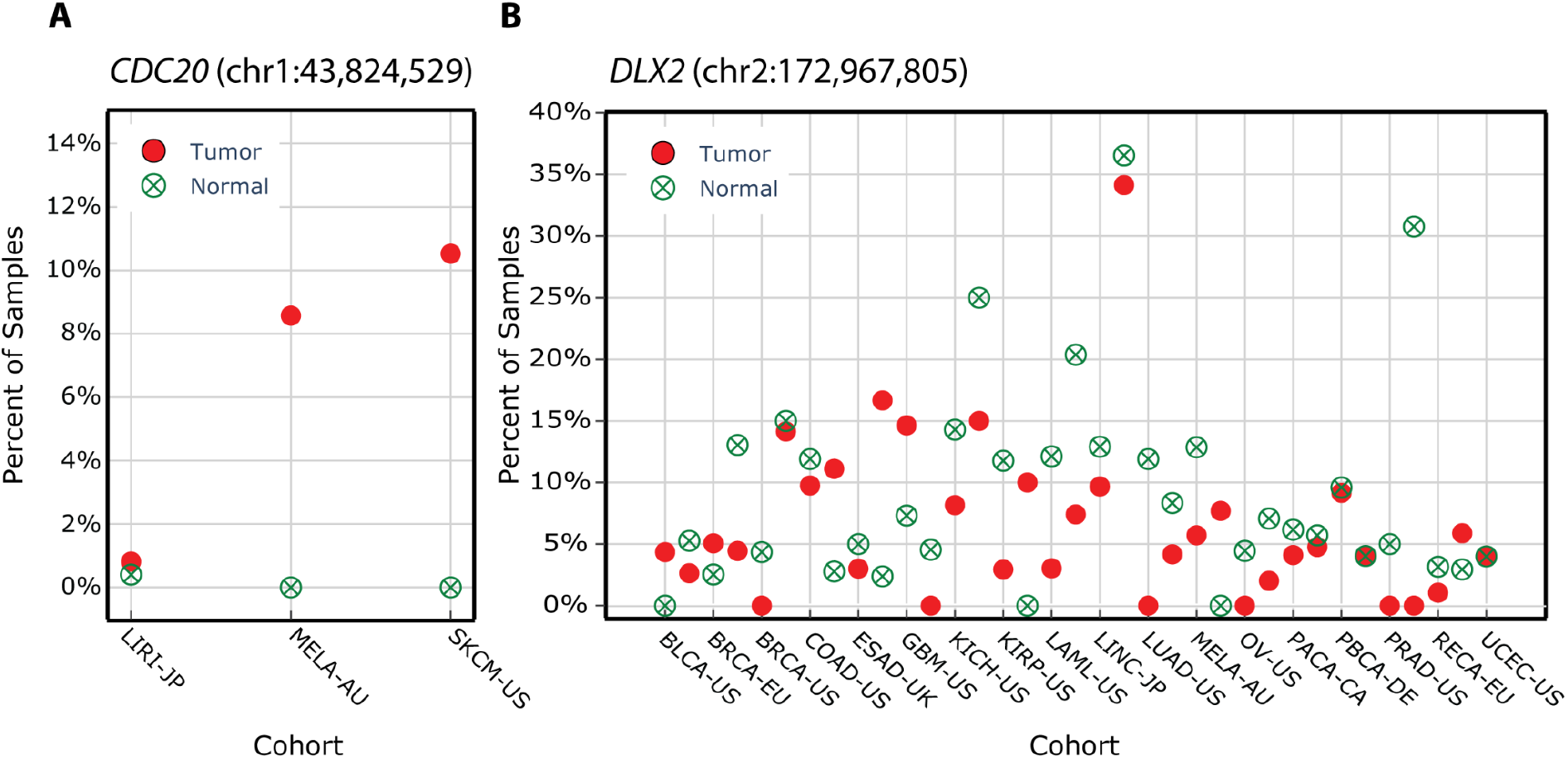
Two examples of a DeepPileup plot of the percentage of samples with a minor allele frequency (MAF) above 25% for both tumor and normal samples per cohort. (A) A technically valid genomic position due to having almost no signal within normal samples but signal within tumor samples (b) A genomic position that could be considered to be noisy due to observing signal in both the control and tumor in a majority of cohorts.

## Notes

### Competing Interest Statement

The authors have declared no competing interest.

